# IMPACT OF PHENOLIC ACIDS ON THE ENERGY METABOLISM AND LONGEVITY IN *C. ELEGANS*

**DOI:** 10.1101/2020.06.23.166314

**Authors:** Benjamin Dilberger, Selina Weppler, Gunter P Eckert

**Affiliations:** Institute of Nutritional Sciences, Laboratory for Nutrition in Prevention and Therapy, Biomedical Research Center Seltersberg (BFS), Justus Liebig University Giessen, Schubertstrasse 81, 35392 Giessen, Germany

**Keywords:** *C. elegans*, PX627, polyphenols, metabolites, hormesis, mitochondria

## Abstract

**Introduction:** Aging represents one of the major risk factors for metabolic diseases, such as diabetes and obesity, or neurodegeneration. Polyphenols and its metabolites, especially simple phenolic acids, have gained more and more attention as a preventive strategy for age-related, non-communicable diseases, due to their hormetic potential. Using the nematode *Caenorhabditis elegans* (*C. elegans*) we investigate the effect of protocatechuic, gallic and vanillic acid to improve mitochondrial function and health associated parameters as a preventive measure.

**Methods:** Lifespan, heat-stress resistance and chemotaxis of *C. elegans* strain PX627, as a specific model for aging, were assessed in 2-day and 10-day old nematodes. Mitochondrial membrane potential (ΔΨm) and ATP generation of young and aged nematodes were measured. mRNA expression levels of longevity and energy metabolism-related genes were determined using qRT-PCR.

**Results:** All phenolic acids were able to significantly increase the nematodes lifespan, heat-stress resistance and chemotaxis at micromolar concentrations. While ΔΨm was only affected by age, vanillic acid significantly decreased ATP concentrations in aged nematodes. Genetic analysis revealed increased glycolytic activity mediated through vanillic acid, suggesting improved thermogenesis.

**Conclusion:** While life- and health-span parameters are positively affected by the investigated phenolic acids, the concentrations applied were unable to impact mitochondrial performance, suggesting hormesis. In contrast to the other phenolic acids, vanillic acid showed potential in regulating glucose homeostasis, making it a prime candidate for future diabetes and obesity focused approaches.

## Introduction

Aging represents a major risk factor for the development of cardiovascular diseases, neurodegenerative diseases, as for example Alzheimer’s or Parkinson’s [1], or metabolic disorders, such as type II diabetes and obesity [2]. Over the last decades, growing evidence suggests mitochondrial function to play a key role during aging [3, 4]. The “mitochondrial theory of aging”, first proposed by Harman in 1956 [5], suggests the accumulation of reactive oxygen species (ROS) to cause oxidative damage, induce mitochondrial dysfunction and thus contribute to aging and the development of related diseases [6, 7]. Although an imbalance of exaggerated ROS are associated with DNA mutations or oxidative damage to lipids and proteins [8], recent evidence suggests mild doses of stressors to be beneficial [9]. Hormesis describes a dose-response phenomenon in which high doses of exposure become harmful, low doses, however, are able to activate protective pathways resulting in a health promotion [10, 11].

Mitochondria represent a unique organelle, conserved over long evolutionary distances and across species, making nematodes an ideal organism to investigate mitochondrial function [6, 12]. According to a recent report mitochondria of nematodes even poses an cold-tolerance mechanism comparable to mammalian non-shivering thermogenesis, allowing mitochondria to generate heat rather than ATP [13].

Polyphenols, as secondary plant metabolites, have emerged as a promising nutritional strategy to prevent oxidative stress and comprise health promoting capacities [14]. Besides their ability to directly scavenge radicals, they gained increasing attention as hormetic agents. Mild concentrations have proven to activate phase II enzymes and thus induce antioxidant defence [15, 16]. Especially treatment of metabolic disorders, for example diabetes or obesity, with polyphenols has gained growing attention [17–19]. However, several clinical trials have failed to provide positive evidence of antioxidant therapies [20]. A possible reason might be the poor bioavailability of polyphenols. An estimate of only 5-10% can be absorbed by the intestine, while the rest is being passed on and metabolized by the gut microbiota [21]. For this reason, especially pre-fermented polyphenolic metabolites, especially simple phenolic acids, gained growing attention, as a strategy to make polyphenols more accessible for absorption [22, 23]. In a recent study, we found the phenolic acid protocatechuic acid (PCA) to be a powerful agent to improve mitochondrial functions and induce longevity [24]. Using the nematode *Caenorhabditis elegans*, we further elucidate the hormetic potential of phenolic acids, metabolites of polyphenolic fermentation, on mitochondrial functions, as well as life- and health-span parameters. After initial screening of four phenolic acid’s three, namely aforementioned PCA, gallic acid (GA) and vanillic acid (VA), were chosen for further investigations (**Graphic 1**).

**Graphic 1:**
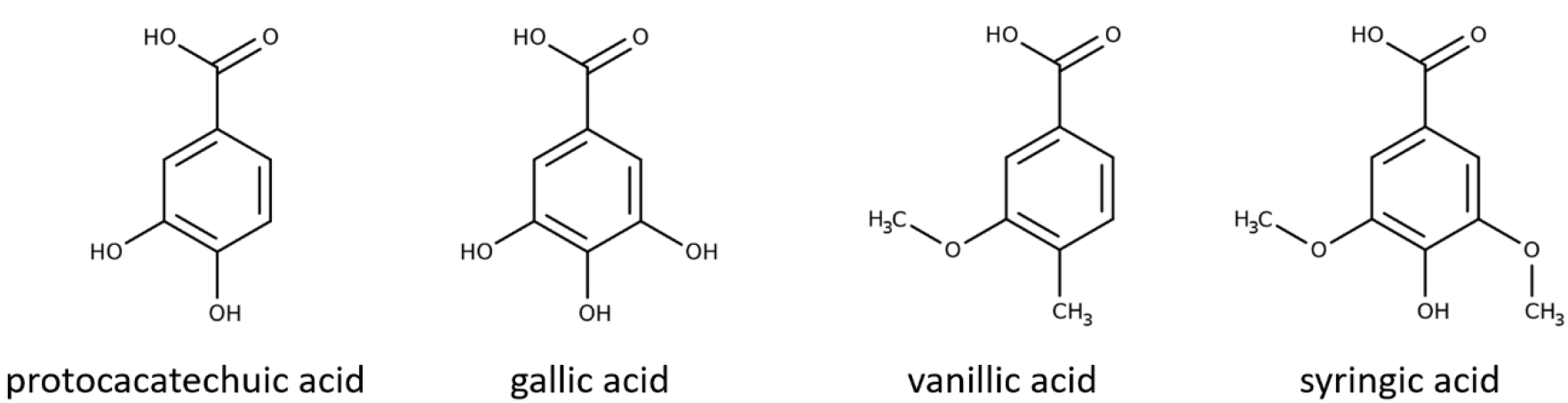
**Chemical structure** of the tested protocatechuic acid, gallic acid, vanillic acid and syringic acid.

We recently established the transgenic strain PX627, allowing the cultivation of synchronous populations free from progeny, without the of use 5-fluoro-2’-deoxyuridine (FUdR), which is known to affect mitochondrial function and longevity [25, 26]. We compare the effects of different phenolic acids in young and aged populations, regarding their impact on mitochondrial functions, as well as glycolytic involvement in energy metabolism and how it impacts the nematodes fitness and survival.

## Material and Methods

### Chemicals

Purity of the chemicals used were of the highest order from Sigma (St. Louis, MO, USA) or Merck (Darmstadt, Germany).

### Nematode and bacterial strains

*C. elegans* strain PX627 (fxls1 I; *spe-44(fx110*[spe-44::degron])IV) was ordered from the *Caenorhabditis Genetics Centre* (University of Minnesota, MN, US). PX627 represents a modified wild-type strain with an inserted auxin-inducible degradation system targeting the *spe-44* gene mediating sterility, backcrossed 5 times [27]. Nematodes were maintained on nematode growth medium (NGM) agar plates seeded with *E. coli* OP50 at 20°C according to standard protocols [28]. Synchronous populations were generated for all experiments with a standard bleaching protocol [29].

### Cultivation and Treatment

Synchronous larvae were counted, adjusted to 10 larvae per 10 μl and raised in cell culture flasks (Sarstedt, Nürmbrecht, Germany) or OP50 spread NGM plates. Liquid OP50-NGM was given as a standardized food source with a volume 4.4-fold of the larvae containing volume. Nematodes were maintained under continuous shaking at 20°C until they reached adulthood, which was defined as day-0. Treatments with protocatechuic acid (PCA), gallic acid (GA), vanillic acid (VA), syringic acid (Syr) and 1% ethanol-M9 solution as control were administered at day-0. Measurements were performed at day-2 (d2) and -10 (d10). To ensure food supply *ad libitum*, for old groups OP50-NGM and effectors were discarded and renewed at day-2 and -6.

### Lifespan assay

After completing the L4 larval stage 60 healthy animals per group were transferred onto fresh NGM E. coli containing plates using a sterilized platinum wire. Effectors were incorporated into the OP50 culture with the concentration as given. To ensure sufficient food supply and to separate dead from alive nematodes, they were transferred onto fresh effector containing agar plates every two days. Nematodes that crawled off the plates were censored from the experiment. Lifespan curves were statistically compared using the log-rank test.

### Heat-stress resistance assay

The nematodes ability to tolerate heat-stress resistance at 37 °C was assessed using a microplate assay. Nematodes were thoroughly washed with M9-Tween® 20 buffer to discard residual bacteria and individually separated into each well of a black 384-well low-volume microtiter plate (Greiner Bio-One, Frickenhausen, Germany), prefilled with 6.5 μl M9-buffer/Tween® 20 (1% v/v). A volume of 7.5 μl SYTOX™ green (final concentration 1 μM; Life Technologies, Karlsruhe, Germany) was added for fluorescent detection. The fluorophor penetrates only into cells with compromised plasma membrane and gets fluorescent through DNA intercalation. Plates were sealed with a Rotilab sealing film (Greiner Bio-One, Frickenhausen, Germany) to prevent water evaporation. Heat-shock (37°C) was applied and fluorescence measured with a ClarioStar Platereader (BMG, Ortenberg, Germany) every 30 min over the course of 17 h. The excitation wavelength was 485 nm and the emission 538 nm.

### Chemotaxis assay

Agar plates were divided into four quadrants. Sodium azid (0.5 M) was mixed in same parts with ethanol (95%) as control, or diacetyl (0.5%) as attractant. A volume of 2 μl for either control or attractant were added to the centre of two opposite quadrants. Approximately 100 animals were placed in the centre of each plate and checked after 1 h, to which quadrant they crawled to. Chemotaxis index calculated ((number of attractant – number of control) / number total) [30, 31].

### Isolation of mitochondria

A population of 5,000 gravid adults were transferred into ice-cold isolation buffer (300 mM Sucrose, 5 mM TES, 200 μM EGTA, pH 7.2). A Balch homogenizer (Isobiotec, Heidelberg, Germany) was used to generate a sufficient and mitochondria enriched fraction. In short, nematodes were passed through the homogenizer’s chamber for five times, with a 12μM clearance, using a 1 ml glass syringes (SGE Syringe, Trajan, Australia). To clear the homogenate from debris and larger worm fragments it was centrifuged at 800 g for 5 minutes at 4°C (Heraeus Fresco 21, Thermo Scientific, Langenselbold, Germany) and the supernatant collected. Mitochondria were concentrated with an additional centrifugation at 9,000 g for 10 minutes at 4 °C. The crude mitochondria containing pellet was resuspended in 70 μl swelling buffer (SWB) (0.2 M sucrose, 10 mM MOPS-Tris, 5 mM succinat, 1 mM H_3_PO_4_, 10 μM EGTA, 2 μM rotenone). Aliquots were shock frozen in liquid nitrogen for determination of protein content.

### Mitochondrial membrane potential (ΔΨm)

Rhodamine 123 (Rh123) was used to assess the mitochondrial membrane potential in 25 μl of isolated mitochondria. Fluorescence was detected in a black 96 well-plate with a ClarioStar Platereader (BMG, Ortenberg, Germany). To ensure mitochondrial integrity ΔΨm was measured for 30 minutes and after reaching equilibrium, 500 nM FCCP was added to evaluate the ΔΨm dependent effect on the quenching of Rh123. Results were normalized to protein content.

### Nematode Homogenization

Adult nematodes were collected, washed, shock frozen and boiled for 15 minutes, to denaturate degrading enzymes. Supernatants were collected, after centrifugation at 15,000 g for 10 minutes. The supernatant was divided into aliquots for ATP, protein, lactate and pyruvate measurements. Lactate and Pyruvate aliquots were stored at −80°C, whereas ATP and protein concentrations were assessed immediately.

### ATP measurement

ATP luminescence was determined in triplicates using the ATPlite luminescence assay (Perkin Elmer, Waltham, MA, USA) with a ClarioStar Platereader (BMG, Ortenberg, Germany). Results were normalized to protein concentrations.

### Protein Quantification

To determine protein concentrations the Pierce™ BCA Protein Assay Kit (Thermo Fisher Scientific, Waltham, MA, USA) was used, according to the manufacturer’s guidelines, with bovine serum albumin as a standard.

### Quantitative real-time PCR

The RNeasy Mini Kit (Qiagen, Hilden, Germany) was used to isolate total RNA, according to the manufacturer’s guidelines. Complementary DNA was synthesized from 1 μg total RNA with an iScript cDNA Synthesis Kit (Bio-Rad, Munich, Germany). A CFX 96 Connect™ system (Bio-Rad, Munich, Germany) was used to perform qRT-PCR experiments with primers purchased from Biomers (Ulm, Germany). Oligonucleotide primer sequences, primer concentrations and product sizes are listed in **Table 1**. Gene expression levels were normalized to amanitin resistant (*ama-1*) and actin (*act-2*). According to the MIQUE guidelines a melting curve was prepared for each primer pair resulting in one significant melting peak.

**Table 1:**
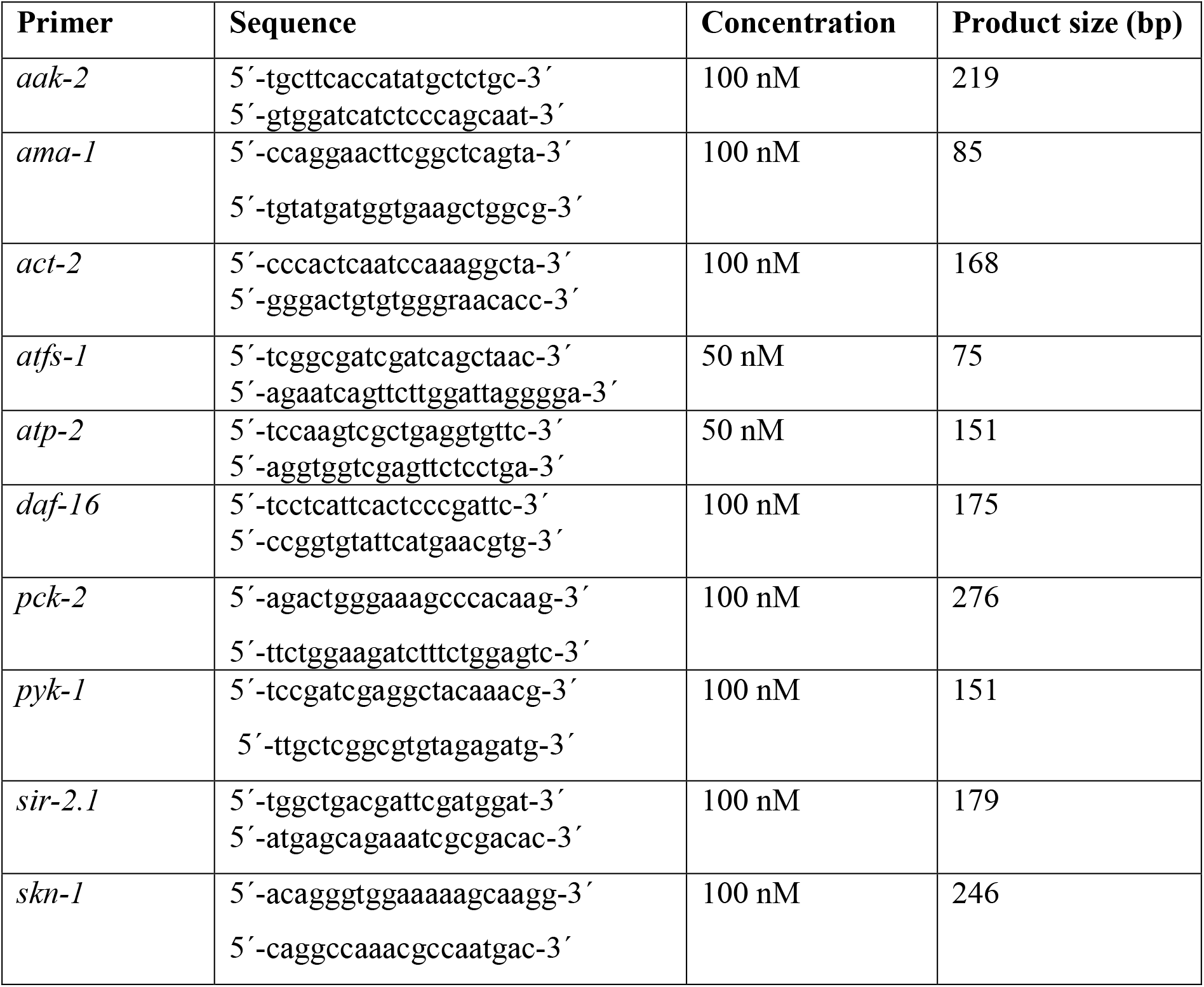
Oligonucleotide primer sequences and product sizes for quantitative real-time PCR. bp: base pairs.

**Table 2:** Relative normalized mRNA expression levels of 2- and 10-day old PX627 nematodes treated with control, 100 μM PCA, 100 μM GA or 250 μM VA. mRNA expression of 2-day control is 100%. n = 7-10; mean ± SEM; one-way ANOVA with Tukey’s multiple post-test. * = compared to 2d group of the same treatment; b = compared to 10d control group. Results are normalized to mRNA expression of *amanitin resistant-1* (*ama-1*) and *actin-2* (*act-2*).

| | control | | PCA 100 μM | | GA 100 μM | | VA 250 μM | |
| --- | --- | --- | --- | --- | --- | --- | --- | --- |
|  | 2d | 10d | 2d | 10d | 2d | 10d | 2d | 10d |
| *daf-16* | 100 ± 8.7 | 138.9 ± 23.6 | 110.1 ± 6.4 | 101.4 ± 12.4 | 93.4 ± 10.4 | 75.6 ± 15.6<br>$p^b = 0.033$ | 112.1 ± 10.5 | 93 ± 14.5 |
| *sir-2.1* | 100 ± 11.2 | 187.7 ± 59.4 | 115.6 ± 24.2 | 202 ± 24.2 | 113 ± 20.8 | 259.2 ± 69.1 | 134.5 ± 19.7 | 540.1 ± 161.1<br>$p^{**} = 0.004$<br>$p^b = 0.036$ |
| *aak-2* | 100 ± 15 | 79.7 ± 20.5 | 122.1 ± 19.1 | 53.2 ± 5.8<br>$p^* = 0.017$ | 90.1 ± 9.7 | 75.5 ± 14.2 | 79 ± 12.4 | 35.6 ± 7.9 |
| *skn-1* | 100 ± 16.9 | 154.1 ± 31.2 | 166.4 ± 25.1 | 118.9 ± 18.9 | 111.4 ± 16.4 | 99.6 ± 15.7 | 135.5 ± 16.5 | 193.1 ± 27.7 |
| *atp-2* | 100 ± 11.7 | 121.4 ± 19.6 | 108.5 ± 11 | 121.5 ± 28.8 | 110.1 ± 15.7 | 72.4 ± 12.9 | 120.9 ± 19.3 | 80.4 ± 12.6 |
| *atfs-1* | 100 ± 14.5 | 157.5 ± 38.7 | 124.5 ± 14.3 | 108.2 ± 26.4 | 137.6 ± 21.3 | 74.3 ± 9.6 | 144.2 ± 19.9 | 359.2 ± 94.7<br>$p^{**} = 0.003$<br>$p^{bb} = 0.009$ |
| *pyk-1* | 100 ± 18 | 86.9 ± 17 | 80.7 ± 8 | 104.4 ± 41.1 | 99.3 ± 18.2 | 104.4 ± 16.8 | 127.8 ± 20.2 | 334.3 ± 67<br>$p^{***} < 0.001$<br>$p^{bbb} < 0.001$ |
| *pck-2* | 100 ± 9.9 | 91.2 ± 28.1 | 91.4 ± 5.2 | 56.5 ± 12 | 67.8 ± 6.9 | 56.9 ± 8 | 53 ± 8.5 | 39.7 ± 6.3<br>$p^b = 0.049$ |

### Statistics

Unless otherwise stated, values are presented as mean ± standard deviation (SD). Statistical analyses were performed by applying a one-way analysis of variance (ANOVA) with Tukey’s multiple comparison *post-test* (Prism 8.3 GraphPad Software, San Diego, CA, USA). Statistical significance was defined for *p* values *p* *^/a/b^ < 0.05, *p*\*\*^/aa/bb^ < 0.01 and *p*\*\*\*^/aaa/bbb^ < 0.001. (*) is used to differentiate between groups of the same treatment, whereas (a) compares 2d treatment groups to 2d controls and (b) 10d treatment groups to 10d controls.

## Results

### Life- and Health-span assessment

To investigate the potential effects of the chosen phenolic acids protocatechuic acid (PCA), gallic acid (GA), vanillic acid (VA) and syringic acid (Syr), it was first necessary to evaluate a target concentration for further investigation. For practical reasons a heat-stress resistance assay was chosen to elucidate the nematodes survival after treatment with all substances at concentrations of 100 μM, 250 μM and 500 μM, respectively (**Fig.1**).

**Fig. 1.**
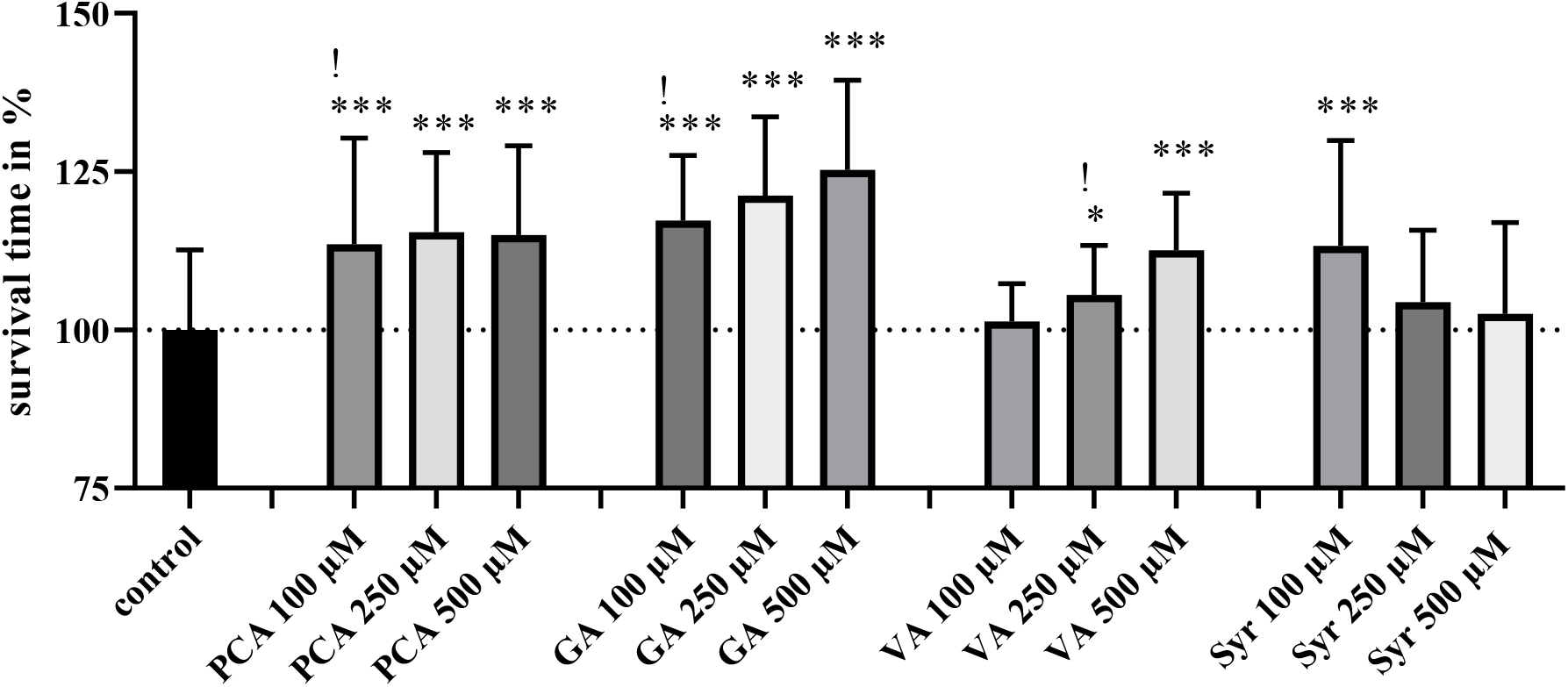
Concentration evaluation via heat-stress: Assessment of heat-stress resistance at 37°C for PCA, GA, VA, and Syr, with concentrations of 100 μM, 250 μM and 500 μM, respectively. Heat-stress resistances are shown in percent, compared to the control group set to 100%. Target concentrations of the chosen metabolites to further investigate are marked with a “!”. n > 57; log-rank (Mantel-cox) test; p* < 0.05 and p*** < 0.001.

**Fig. 2.**
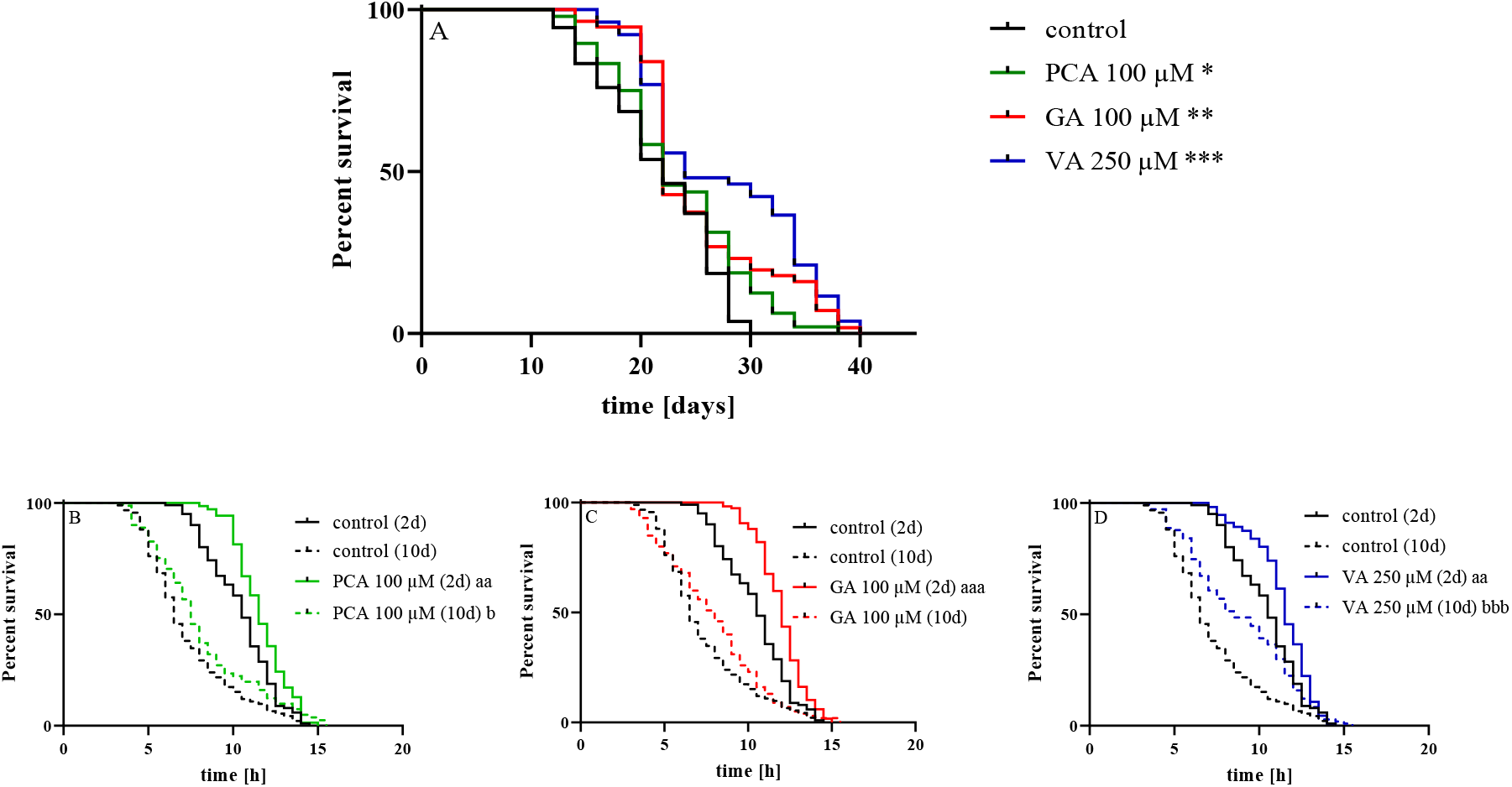
Survival and stress-resistance: The survival under physiological conditions was determined for the same chosen treatments **(A)**. Stress resistance at 37°C was assessed in young (2d) and aged (10d) PX627 after treatment with 100 μM PCA **(B)**, 100 μM GA **(C)** and 250 μM VA **(D)**. Significant differences between 2d groups are indicated with “a” and those of 10d groups using; for stress-resistance “b”. n > 54 and for survival n > 48; log-rank (Mantel-cox) test; p^a/b^ < 0.05, p^aa/bb^ < 0.01 and p^aaa/bbb^ < 0.001.

**Fig. 3.**
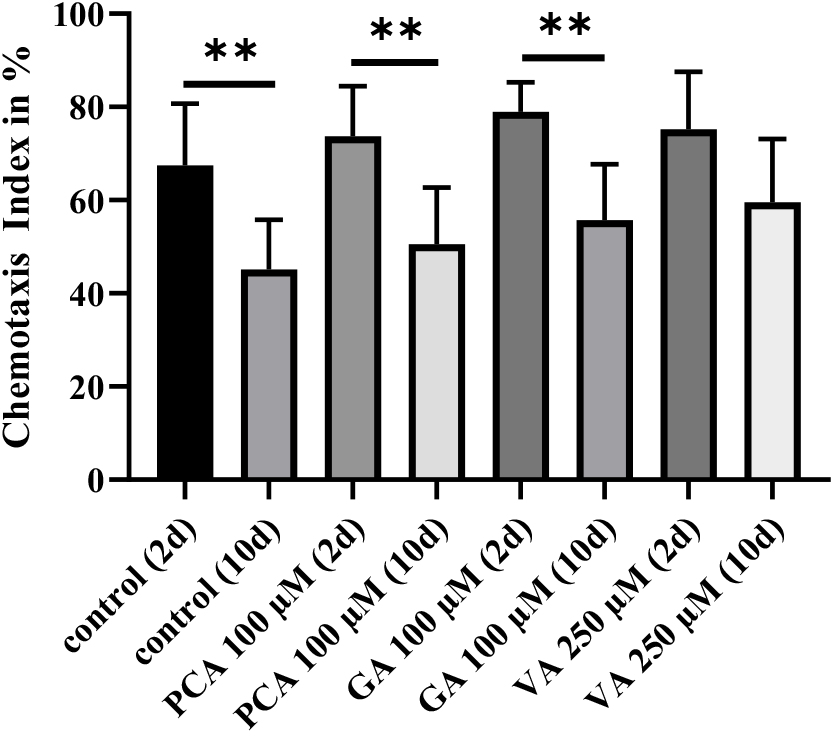
Chemotaxis: Chemotaxis index in percent of 2d and 10d old nematodes treated with 100 μM PCA, 100 μM GA and 250 μM VA; n = 8; mean ± SD; one-way ANOVA with Tukey’s multiple comparison post-test; p** < 0.01.

**Fig. 4.**
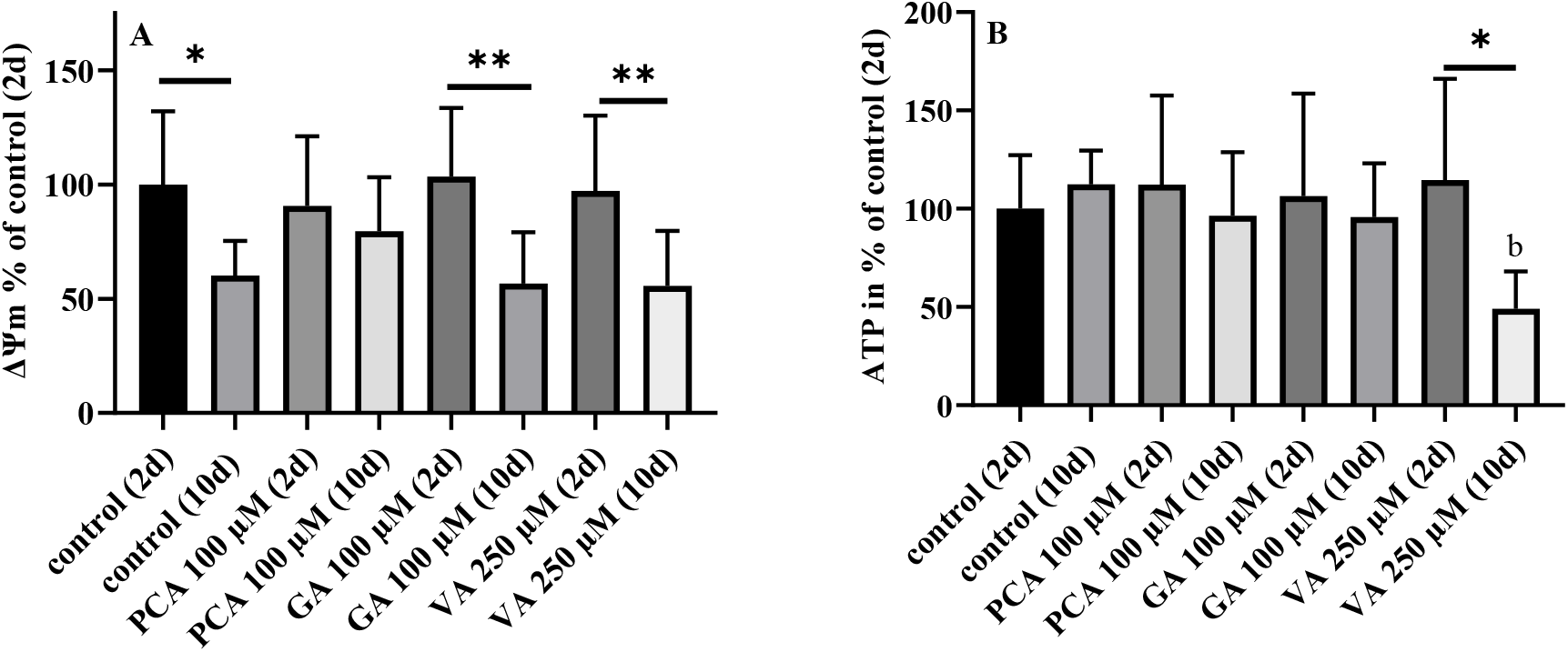
Mitochondrial Parameters: Assessment of (**A**) mitochondrial membrane potential (ΔΨm) and (**B**) ATP concentrations of 2d and 10d old nematodes treated with 100 μM PCA, 100 μM GA and 250 μM VA. n = 9-13; mean ± SD; one-way ANOVA with Tukey’s multiple comparison post-test; p*^/b^ < 0.05.

The metabolites PCA, GA and VA, with the lowest concentration reaching a level of significance, were chosen for further investigation. Syringic acid was disregarded, since its optimal concentration appears to fall below the assessed range (for further explanation see Discussion).

To confirm a beneficial effect of the chosen metabolites and concentrations (100 μM PCA, 100 μM GA and 250 μM VA), the nematodes survival was determined under physiological conditions at 20°C. While all treatment groups experienced a significant increase in longevity, the effect was most potent for VA. Observation of young and aged nematodes capability to tolerate heat-stress showed similar results. Both age-stages of all treatments were more resilient to thermal stress. While the effect was comparable in both age-stages for PCA, GA appears to impact especially young and VA especially aged nematodes.

Analysis of the animal’s chemotactic ability to locate food reflects the findings of the previously conducted lifespan and heat-stress assessment. The chemotaxis index of all groups were numerically increased, compared to their respective young and aged control groups. In contrast to PCA and GA, treatment with VA, however, increased the chemotaxis index of 10-day old nematodes to an extent being no longer significantly decreased compared to 2-day old VA treated nematodes.

### Mitochondrial parameter

Accelerated age significantly reduces the mitochondrial membrane potential (ΔΨm). The ΔΨm represents the driving force for ATP generation [32]. ATP concentrations were, regardless of the reduced membrane potential, unaffected in aged nematodes. Whereas, concerning ΔΨm, treatment with GA and VA reflects the findings of their respective controls, the ΔΨm of aged PCA-treated nematodes was slightly, although not significantly, improved to a level being no longer different from the young PCA-treated group. ATP concentrations were affected neither by age nor treatment. The exception of aged VA-treated nematodes, whose concentrations were significantly reduced compared to 2d VA-treated nematodes and 10d controls, even though the ΔΨm was not affected.

### qRT-PCR

Analysis of mRNA expression reveals a significantly decreased expression of *aak-2* in aged PCA treated nematodes, as well as a mentionable, but not significant, increase in *sir-2.1* expression, compared to aged untreated controls. VA treatment significantly increased longevity-related marker genes *sir-2.1* as well as numerically *daf-16* and *skn-1* in aged nematodes, compared to their respective aged controls. Expression of UPR^mt^ marker *atfs-1*, was significantly increased in aged VA treated nematodes to young VA treated nematodes and untreated aged controls. Furthermore, VA-treatment significantly up-regulated *pyk-1*, which encodes for the enzyme pyruvate kinase, responsible for glycolytic ATP production, in aged nematodes compared to their young counterparts and respective controls. In contrast, *pck-2*, as a marker gene for gluconeogenesis, was significantly down-regulated in aged VA-treated nematodes compared to untreated controls.

## Discussion

Increasing evidence suggests a close relationship between mitochondrial function and longevity [6, 33]. During oxidative phosphorylation electrons are able escape mitochondrial respiration, forming reactive oxygen species (ROS), which can cause cellular damage in form of lipid and DNA oxidation [34, 35]. Mitochondrial dysfunction, associated with increased ROS production, is reported to increase during aging, causing a further increase in oxidative stress. This results in a vicious cycle of increased ROS through age-associated mitochondrial dysfunction, which again accelerates the aging process [36]. Polyphenols, due to their potential in ROS-scavenging, have emerged as a potentially powerful strategy to ameliorate oxidative stress and thus induce health and longevity [37–39].

However, bioavailability of complex polyphenols is supposed to be low and they are metabolized by intestinal microbiota, into smaller intermediates, before they can be taken up [23]. Gallic acid, for example, a simple benzoic acid, is attributed with a good absorbability [22]. In general, phenolic acids represent a subclass of polyphenols attributed with, for example, anti-oxidative, anti-inflammatory, or anti-cancer properties. In fact, the antioxidant capacities of some phenolic acids were found to exceed that of well-known antioxidant vitamins [22]. In a previous study we demonstrated beneficial effects for the phenolic acid PCA on mitochondrial function and longevity in the nematode model *C. elegans* [24]. Here we follow up on the health promoting capacities of PCA using a recently described nematode model enabling the comparison of young and aged nematodes without the use of 5-fluoro-2’-deoxyuridine, which heavily affects mitochondrial function, to maintain synchronous populations [26]. In addition, vanillic acid, a precursor metabolite of PCA [40], as well as syringic acid (Syr) and its metabolite gallic acid (GA) [41], of which plasma concentrations were found to be increased after administration [42–45], were co-assessed regarding their effect on the energy metabolism of young and aged nematodes, due to their structural similarities [46].

A heat-stress resistance assay was used to determine a target concentration for each phenolic acid in young nematodes. The effect of three concentrations (namely 100 μM, 250 μM and 500 μM), covering a range commonly used for phenolic investigations in *C. elegans* [24, 47, 48], was assessed and the lowest concentration reaching a level of significance (*p*\* < 0.05) chosen for further investigation.

### Life- and health-span assessment

Target concentrations of the remaining three phenolic acids were evaluated regarding their potential on the nematodes lifespan under physiological conditions to validate health-promoting capacities. In fact, all substances significantly increased the nematodes median survival, with vanillic acid being the most potent. A lifespan extension for all three phenolic acids is well documented [24, 45, 47, 49]. Protection against thermal stress has already been demonstrated during determination of target concentrations in young, but not in aged 10-day old, nematodes. Again, all three substances could significantly improve the nematodes ability to tolerate thermal stress. In young nematodes the most distinct protection was conveyed by gallic acid, but in aged nematodes this was the case for vanillic acid. This is in accordance with Saul et al., who reported gallic acid to prolong the nematodes lifespan and to protect against thermal stress [45]. While PCA has been extensively investigated and its health promoting capacities are well described [24, 47], analysis of VA on the nematodes health (and mitochondrial function) is novel to the literature. To further strengthen these findings we analysed key genetic marker genes for longevity and stress-resistance *daf-16* and *sir-2.1* [50, 51]. While *daf-16* was mostly not affected, *sir-2.1* expression was increased after by all, but especially VA, treatment. This suggests the life- and health-span promotion to be induced by sirtuins. Previously we reported *daf-16* and *sir-2.1* to be significantly increased after PCA administration, however it is to mention concentrations were almost 8-times higher compared to this approach [24].

Chemotaxis, the nematodes ability to locate food, a marker for neurological function and acceleration of aging [30, 31], reflects the findings of our lifespan and thermo-tolerance experiments. All treatment groups showed numerically increased chemotaxis indices, compared to their respective controls, as well as an age-dependent decline within the same treatment group. Again, especially VA protected against loss in neuronal function. VA notably elevated the chemotaxis index of aged nematodes to a level being no longer significantly decreased compared to their young treated counterparts. Taken together, this suggests all three phenolic acids to be beneficial on health-associated parameters, with an emphasis on vanillic acid, which appears to develop its full potential in aged specimens.

For this reason we argue, although a beneficial effect of all three phenolic acids on life- and health-span parameters could be measured, with the exception of VA, the concentrations administered were too low to significantly impact genetic markers, suggesting a hormetic mode of action [52].

### Mitochondrial involvement

Mitochondria are responsible for the majority of the cells energy production in form of ATP. Approximately 90% of ATP are generated by the mitochondrial respiration chain, which depends on an electron gradient across the inner mitochondrial membrane, referred to as mitochondrial membrane potential (ΔΨm) [32, 53, 54]. ΔΨm is known to decline with age [26, 55], which can be further evidenced by our results. However, neither phenolic acid was able to improve ΔΨm compared to their respective control group, suggesting no mitochondrial modulation by either substance at the concentration administered. Expression of *skn-1*, ortholog of mammalian PGC1α and responsible for mitochondrial biogenesis [56, 57], reflects these findings. However, concerning polyphenols the literature are more contradicting. Aitken et al. reported adverse effects of various polyphenols, but mostly catechins, on the ΔΨm in human spermatozoa [58]. Rafiei et al. and Li et al. demonstrated improved mitochondrial function after administration of prominent polyphenols, as for example resveratrol and quercetin, in HepG2 and tea polyphenols in VEC cells [59, 60]. However, it is to mention that, also belonging to the family of polyphenols, these substances vary in structure and their bioactivity from the here presented simple phenolic acids [22]. Previously we reported a significantly improved mitochondrial membrane potential after 780 μM PCA administration [24]. Although a numerical up-regulation could be seen, due to lesser concentrations used we were unable to reproduce similar effects in this study. Since the other phenolic acids were also unable to significantly improve mitochondrial performance, we argue although a beneficial effect cannot be ruled out, the concentrations applied were too low to significantly improve the nematodes’ ΔΨm. This is in support of our hypothesis of a hormetic effect of the investigated phenolic acids, since even though mitochondrial function appears mostly unaltered, longevity was induced. This appears feasible since a dose-depended effect of PCA to ameliorate rotenone induced toxicity has been shown in PC12 cells [61]. Interestingly, VA significantly up-regulated the expression of *atfs-1*, indicating an increased mitochondrial unfolded protein response, generally associated with increased longevity and age [62, 63], as can be further evidenced by our life- and health-span assessment.

Although ΔΨm was unaffected by the administered phenolic acids and dropped only during senescence, ATP concentrations were stable throughout all treatment groups and age-stages, with the exception of aged VA-treated nematodes. Unaltered expression levels of *atp-2*, a subunit of complex V, supports these findings. Generally this is in line with previous findings, suggesting organisms maintain stable ATP concentrations in respect to their needs [26, 64, 65]. This is accomplished on the cost of increased ROS generation, due to the cells inability to maintain their ΔΨm [26, 66]. Glycolytic activity was assessed, as a second energy-generating pathway [67], to explain the discrepancy of ATP concentrations in aged VA-treated nematodes. Pyruvate kinase (PK), a key enzyme of glycolysis and glucose homeostasis, transfers a phenolic group from phosphoenolpyruvate onto ADP, thus generating ATP and pyruvate [68]. However, expression of *pyk-1,* encoding for PK, was significantly up-regulated, suggesting an increased rather than decreased glycolytic activity. Considering the decreased *pck-2* expression, a genetic marker for gluconeogenesis, increased glycolysis due to VA administration becomes even more evident. Glycolysis and gluconeogenesis are generally considered as opposing metabolic pathways [69], thus ruling glycolysis out to be responsible for the observed ATP-depletion. Decreased gluconeogenesis mediated by vanillic acid has already been demonstrated [70], as well as for a VA-containing extract [71], thus strengthening our findings. The ATP-depletion in aged VA-treated nematodes could be explained by increased mechanisms similar to thermogenesis. Just as brown adipose tissue in mammals, nematodes are also able to respond to cold temperatures via activation of the cAMP-PKA, thus increasing their fat utilization [13]. This is in accordance with Han et al. and Jung et al., who previously found vanillic acid to promote thermogenesis in C57BL/6J mice and in primary cultured adipocytes [72, 73]. Mitochondrial autophagy represents another feature commonly associated with increased mitochondrial thermogenesis [74]. Related markers were increased especially for aged VA-treated nematodes, thus further contributing to the body of evidence suggesting vanillic acid to induce thermogenesis.

## Conclusion

In summary, our findings suggest phenolic acids to positively affect the nematodes life- and health-span, possibly via hormesis, since the administered concentrations appear to be too low to directly improve mitochondrial function. Vanillic acid, however, shows promising potential in controlling glucose homeostasis and thermogenesis, hallmarks generally associated with combating type II diabetes or obesity.

## Data Availability

The dataset generated during this study is available from the corresponding author on reasonable request.

